# Genetic liability underlying reward-related comorbidity in psychiatric disorders involves the coincident functions of autism-linked ADGRL1 and hevin

**DOI:** 10.1101/2024.07.03.601736

**Authors:** Kerlys G. Correoso-Braña, Augusto Anesio, Sylvie Dumas, Emmanuel Valjent, Nicolas Heck, Vincent Vialou, Antony A. Boucard

## Abstract

Comorbidity between psychiatric traits is thought to involve overlapping pleiotropic effects from sets of genes. Notably, substance abuse is a shared comorbid condition among various neurodevelopmental disorders with externalizing symptoms such as autism spectrum disorder and attention-deficit hyperactivity disorder, thus hinting at the nucleus accumbens (NAc) as a site for predisposition underlying convergence of genetic influences in reward-related comorbidity. Here, we identify the autism-related gene encoding the adhesion G protein-coupled receptor (aGPCR) Latrophilin-1/ADGRL1 as an essential transducer of reward mechanisms in the NAc. We found that ADGRL1 mRNA is ubiquitously expressed throughout major NAc neuronal populations in mice. A mouse model of pan-neuronal Adgrl1 deficiency in the NAc displayed cocaine-seeking impairments in adult individuals denoting its role in drug-induced reinforcement and reward. Connecting molecular pathways of cocaine-induced learning, we uncover that ADGRL1 constitutes a functional receptor for autism-related cocaine effector molecule hevin/SPARCL1. Indeed, hevin interacts with membrane-expressed ADGRL1 and induces its internalization while stabilizing its uncleaved fraction. Moreover, hevin alters the formation of intercellular adhesion contacts mediated by ADGRL1 and Neurexin-1. Importantly, the functional constitutive coupling between ADGRL1 and various G protein pathways is selectively modulated by hevin stimulation with a bias toward Gi3, Gs, and G13 proteins. These findings unveil the dual role of ADGRL1 and hevin as genetic risk factors for both psychiatric disorders and substance abuse to define the molecular etiology of comorbidity.

## INTRODUCTION

Drug addiction exhibits significant comorbidity with neuropsychiatric disorders such as attention-deficit and hyperactivity disorder (ADHD) and autism spectrum disorders (ASD) ^1^ ^2^ ^3^. An increasing number of studies aimed at characterizing animal models of ASD and ADHD revealed striatal dysfunctions akin to those observed in drug addiction models implicating genes that regulate synaptic transmission as well as behavioral plasticity ^4^ ^5^ ^6^. Despite the identification of some genes as risk factors for developing such dysfunctions, little is known about the mechanisms underlying them, nor how some of these genes’ functions might intersect.

Neuropsychiatric disorders possess a strong genetic component often centered on genes known to be essential for neuronal and glial health ^7^ ^8^ ^9^. Recently, loss-of-function variants of the gene encoding the synaptic adhesion molecule Latrophilin-1 (Lphn1/ADGRL1) were reported in a group of heterozygous patients with developmental disorders such as intellectual disability, epilepsy, ADHD and ASD ^10^. ADGRL1 is a G protein-coupled receptor belonging to the Adhesion clade and characterized by a large extracellular region interspersed with protein-protein interaction motifs or adhesion modules located upstream of an autoproteolytic site which generates two non-covalently associated fragments that cooperate to unify receptor functions. Axonal attraction and synapse formation are among the functions mediated by ADGRL1’s adhesion motifs known to stabilize intercellular contacts through heterophilic interactions with its ligands such as the synaptic adhesion molecules neurexins (NRX) ^11^ ^12^. Ligand-induced activation of ADGRL1 leads to a G protein-dependent modulation of cAMP levels as well as actin cytoskeleton remodeling which results in morphological changes in neurons ^13^ ^14^. Thus, ADGRL1 has the ability to exert pleiotropic effects by integrating extracellular synaptogenic cues into intracellular signaling and cytoarchitectural changes relevant to a range of synaptogenic events.

In recent years, several astrocyte-secreted factors responsible for synaptogenesis have been identified and characterized in neuropsychiatric disorders such as ADHD ^15^, depression ^16^, autism ^17^, schizophrenia ^18^ and substance abuse ^19^ ^20^. Amongst them, hevin also known as SPARCL1 was shown to promote synaptogenesis and synaptic transmission at excitatory synapses in neuronal cultures ^21^. During development, hevin is involved in cortical plasticity ^22^ and thalamocortical connectivity ^23^. Indeed, developmental functions have been attributed to hevin since loss-of-function mutations in the *Sparcl1* gene have also been associated with a risk for developing ASD ^24^ ^25^ ^26^. However, in contrast to other astrocyte-secreted factors, hevin remains highly expressed in the central nervous system during adulthood ^27^, thus suggesting a role in adult synaptic plasticity. Indeed, we recently highlighted the role of hevin in contributing to adult synaptic plasticity induced by the drug of abuse cocaine within the nucleus accumbens (NAc) ^28^. Here we provide evidence that the function of ADGRL1 in the reward-related nucleus accumbens is required for drug-seeking behavior in adult mice, and identify the cocaine-transducing matricellular protein hevin as a functional ligand that modulates the signaling properties of ADGRL1.

## MATERIALS AND METHODS

### Animals

Experiments were performed following the laws for the use of animals in neuroscientific research (European Committee Council Directive 2010/63/EU). Male C57BL6J mice between 7 and 19 weeks of age were used for this research. All animals were maintained on a 12 h light/dark cycle, provided with food and water ad libitum.

### Mouse brain sectioning

Brains were removed immediately after sacrifice and frozen in isopentane at –35 °C. Coronal sections with a thickness of 16 µm, were obtained using a cryostat and mounted on poly-L-Lysine-coated slides (Thermo Fisher Scientific). The slides were stored at -80 °C until their use for in situ hybridization assays.

### Fluorescence in situ hybridization (FISH)

Cryostat-generated brain sections from 8 and 19-week-old C57BL6J mice were fixed and acetylated with 0.25% acetic anhydride/100 mM triethanolamine pH 8.0 at room temperature (RT). Probe hybridization was conducted in 50% formamide (Eurobio) using 1 µg/ml of RNA probes (Supplementary Table 1) labeled with digoxigenin (DIG) or fluorescein (Fluo). Subsequently, the cryosections were blocked before performing immunodetection with HRP-coupled anti-DIG or anti-Fluo antibodies (1:5000) followed by tyramide-based immunolabelling. Nucleus staining was performed with 4’,6-diamidino-2-phenylindole (DAPI). The samples were read on a NanoZoomer 2.0-HT (Hamamatsu Photonics, Hamamatsu City, Japan), and the images analyzed with the NDP.view 2.7.52 software (Hamamatsu Photonics).

### Design and validation of microRNAs

Sequences previously reported to silence ADGRL1 protein expression ^29^. (Supplementary Table 2) were chosen for miR engineering into pcDNA™6.2-GW/EmGFP following the vendor’s instructions (BLOCK-iT Pol II miR RNAi Expression Vector Kits, Invitrogen). RNA interference efficacy was tested by immunoblotting HEK293 cell lysates which were co-transfected with miR constructs and a plasmid encoding full-length mVenus-tagged ADGRL1. For the generation of adeno-associated virus constructs, two efficient individual miRNAs sequences were PCR-amplified: one for EmGFP and miR1, and a second fragment for miR2. These sequences were then cloned in tandem (HiFi assembly, New England Biolabs) into the pAAV2-hSyn-eGFP viral vector (Addgene) by substituting the eGFP sequence. Adenoviruses were produced at the AAV production facility at Institut de la Vision (Paris). The efficiency of the AAV-induced ADGRL1 knockdown was validated in vivo.

### Stereotactic surgery

Adult C57BL6J mice underwent general anesthesia using a mixture of ketamine xylazine and with lidocaine as local anesthesia at the incision site. Adenovirus was injected into the nucleus accumbens; Bregma coordinates: anteroposterior: +1.45, mediolateral: +1.45, dorsoventral: -4.3, with an angle of 10° from the midline.

### Conditioned place preference (CPP)

CPP was performed in a two-compartment maze displaying distinct visual and tactile cues. The following schedule was observed: Day 1 (Pre-test): mice were allowed to freely explore both compartments for 30 min. Day 2-3 (Conditioning): Mice were injected with saline (morning) or cocaine (10 mg/kg, afternoon) and placed in one specific compartment for 20 min. Assignation of the cocaine-paired compartment was adjusted to balance out any pre-existing compartment bias. Day 4 (Test): animals were allowed to explore both compartments for 30 min. The CPP score was calculated as the time spent in the cocaine-paired compartment on day 4 minus time spent in the same compartment on day 1.

### Cell culture and transfection

HEK293 and HEK293T cells (ATCC) were cultured in Dulbecco’s Modified Eagle Medium (DMEM, Corning) supplemented with 10% fetal bovine serum (FBS, Biowest), 2 mM GlutaMAX^TM^ (Gibco) and 1000 U/ml penicillin-streptomycin (In Vitro), at 37 °C in a 5% CO_2_ atmosphere. Cells were transfected by the polyethylenimine method (PEI, Polysciences) using a 1:3 PEI:DNA ratio (w/w) with plasmids described in the SI.

### Western blot

Transfected cells were solubilized in gel loading buffer and analyzed through electrophoresis before being transferred to nitrocellulose membranes (Merck Millipore). Immunodetection was achieved with the primary antibodies (1:1000 in blocking solution): anti-HA (BioLegend), anti-Flag (Sigma-Aldrich), anti-GFP (NovusBio), anti-α tubulin (Developmental Studies Hybridoma Bank). Fluorescently-labeled secondary antibodies were added (anti-mouse IRDye800CW or anti-rabbit IRDye680RD; 1:10,000 ratio; LI-COR) and signals detected in the 800 and 700 nm channels, respectively. Images were processed with Image Studio 5.2.5 software.

### Cell-surface detection assays

**Detection of cell-surface binding (DOCS)**-Cells were co-transfected with indicated plasmids, reseeded 1-day post-transfection in poly-L-lysine-treated (Sigma-Aldrich) 96-well plates and cultured for an additional day. Cells were fixed on ice with 4% paraformaldehyde (PFA, Electron Microscopy Sciences) and incubated with anti-HA primary antibody (1:1000, BioLegend) followed by HRP-coupled secondary antibody (1:2000, MP Biomedicals).

**Detection of expression at cell-surface (DECS)**-Cells expressing Flag-tagged ADGRL1 or empty vector were reseeded as in the above-mentioned DOCS assay and incubated with the conditioned media of HA-tagged hevin- or empty vector-expressing cells for 2 h at 37 °C. Following a fixation step on ice using 4% PFA, the flag-tagged receptor was immunodetected with anti-flag primary antibody (1:1000) and HRP-coupled secondary antibody (1:2000).

For both methods, HRP substrate 3,3’,5,5’-tetramethylbenzidine (TMB, Thermo Fisher Scientific) was added and the colorimetric reaction stopped with 1 N sulfuric acid. Absorbance was read at 450 nm using Cytation5 plate reader (Biotek).

### Detection of ligand retention in total extracts (DOLR)

Cells transfected with the indicated plasmids were incubated with hevin-containing media for 2 h at 37 °C. Excess HA-tagged hevin was removed using a phosphate buffer saline (PBS w/o MgCl_2_, CaCl_2_; Corning) wash and cells were lysed in SDS gel sample buffer for further immunoblotting analysis using anti-HA antibody. Calcium-dependency assays were conducted in the presence of 10mM EGTA.

### Cell-surface labeling assay

Cells were plated on poly-L-lysine-coated coverslips and co-transfected with HA-tagged hevin plasmid along with the indicated plasmids. 48h later cells were fixed in 4% PFA dissolved in PBS:DMEM (1:1 v/v) to achieve hevin detection. Surface-bound HA-tagged hevin was detected using an anti-HA antibody (1:200) followed by Alexa-568 coupled secondary antibody (1:200). Nucleus staining was performed with DAPI and 0.4 µm sectioned images were captured by confocal microscopy through a 63X objective (TCS SP8, Leica Microsystems). Image analysis was performed with Leica Application Suite Software (3.7.4.23463 version).

### Co-immunoprecipitation (Co-IP)

Cells transfected with Flag-tagged ADGRL1 construct or empty vector were incubated along with hevin-containing media for 2 h at 37 °C. Cells were lysed in IP buffer (25 mM Tris, 150 mM NaCl, 1% Triton X-100, 5% glycerol, pH 7.4) and the cell extracts were loaded onto an Anti-DYDDDDK (Flag) affinity resin (Thermo Fisher Scientific). Bound proteins were eluted after the addition of 100 mM glycine, pH 2.8 and neutralized in 1 M Tris, pH 8.5.

### Cell aggregation assay

Cell aggregation assays were performed as previously described ^12^. Cells co-expressing ADGRL1 and GFP (or GFP-only as control) were co-cultured with NRX1β(-SS4) and DsRed co-expressing cells (or DsRed-only as control) in the presence of hevin-containing media or control media under gentle rotation for 90 and 120 min. Samples were visualized by epifluorescence microscopy using a 10X objective (DM IL LED, Leica Microsystems). Image analysis was performed with Leica Application Suite Software (3.7.4.23463 version) and Image J 1.53t. The aggregation index was expressed as: (Σ area of particles above threshold/ Σ area of all particles)*100, threshold set as highest particle area in GFP-only/ DsRed only condition.

### Bioluminescence resonance energy transfer (BRET) assays

Cells (3.5 x 10^5^) were co-transfected with increasing amounts of ADGRL1 to detect receptor-mediated constitutive activity along with plasmids for BRET-based G protein biosensors (TRUPATH) ^30^; consisting of the trimer Gα, β and γ, where Gα is coupled to Rluc8 and Gγ to GFP2. For each condition, the total amount of transfected DNA was adjusted to 1 µg with empty vector and cells were cultured for 48 h in a solid-white plate. Coelenterazine 400a (5uM, GoldBIO.com) diluted in BRET buffer (In mM: 10 HEPES, 1 CaCl_2_, 0.5 MgCl_2_, 4.2 KCl, 146 NaCl, 5.5 glucose, pH 7.4) was added to transfected cells and incubated for 5 min at room temperature. Luminescence and fluorescence emission intensities were separately detected using 410/80 nm and 515/30 nm emission filters of a microplate reader (Cytation5, BioTek). The BRET signal was quantified as the BRET^2^ ratio, consisting of: acceptor signal (GFP2)/donor signal (Rluc8).

### Statistical analysis

Data are represented as mean ± standard error of means (SEM) from at least three independent experiments. All statistical analyses were performed in GraphPad Prism version 6.0. These consisted of Student’s t-test and one-way or two-way ANOVA.

## RESULTS

### ADGRL1 is anatomically and functionally related to the reward pathway

The nucleus accumbens is mainly composed of medium spiny neurons (MSNs), expressing either the dopamine D1- or D2-receptor, whose morphological remodeling is a hallmark of drug-induced neuroadaptations and plasticity underlying addiction ^31^. While cocaine-induced structural plasticity in D1-MSNs is long-lasting, the structural changes in D2-MSNs caused by cocaine are more dynamically regulated ^32^. First, we sought to analyze the molecular context of increased propensity to structural plasticity in D2 MSNs by detecting actively transcribed genes in these cells. We focused on genes encoding cell adhesion molecules based on their role as structural stabilizers of synapses ^33^. Expression of transcripts encoding cell adhesion molecules was analyzed using an RNAseq dataset (Gene Expression Omnibus, GSE94145) generated on tagged ribosome-bound mRNAs and the input fractions of NAc extract (including both Core and Shell) of 10-week-old D2-Ribotag ^34^. Amongst the most highly expressed genes were the ones encoding the Amyloid-beta precursor protein APP and the adhesion GPCR ADGRL1/latrophilin-1, each amounting to at least twice the expression of D2-MSN marker *Drd2* (Figure 1a). Both *App* and *Adgrl1* genes were enriched in the D2-MSNs versus in their neighboring cells from the NAc comprising among others D1-MSNs and astrocytes (Figure 1b). Also present in appreciable amounts, albeit lower than *App* and *Adgrl1*, and at various levels of enrichment in D2-MSNs were distinct members of the neurexin and neuroligin families, as well as latrophilin-2 and 3 (*Adgrl2* and *Adgrl3* respectively) (Figure 1a,b).

**FIGURE 1.**
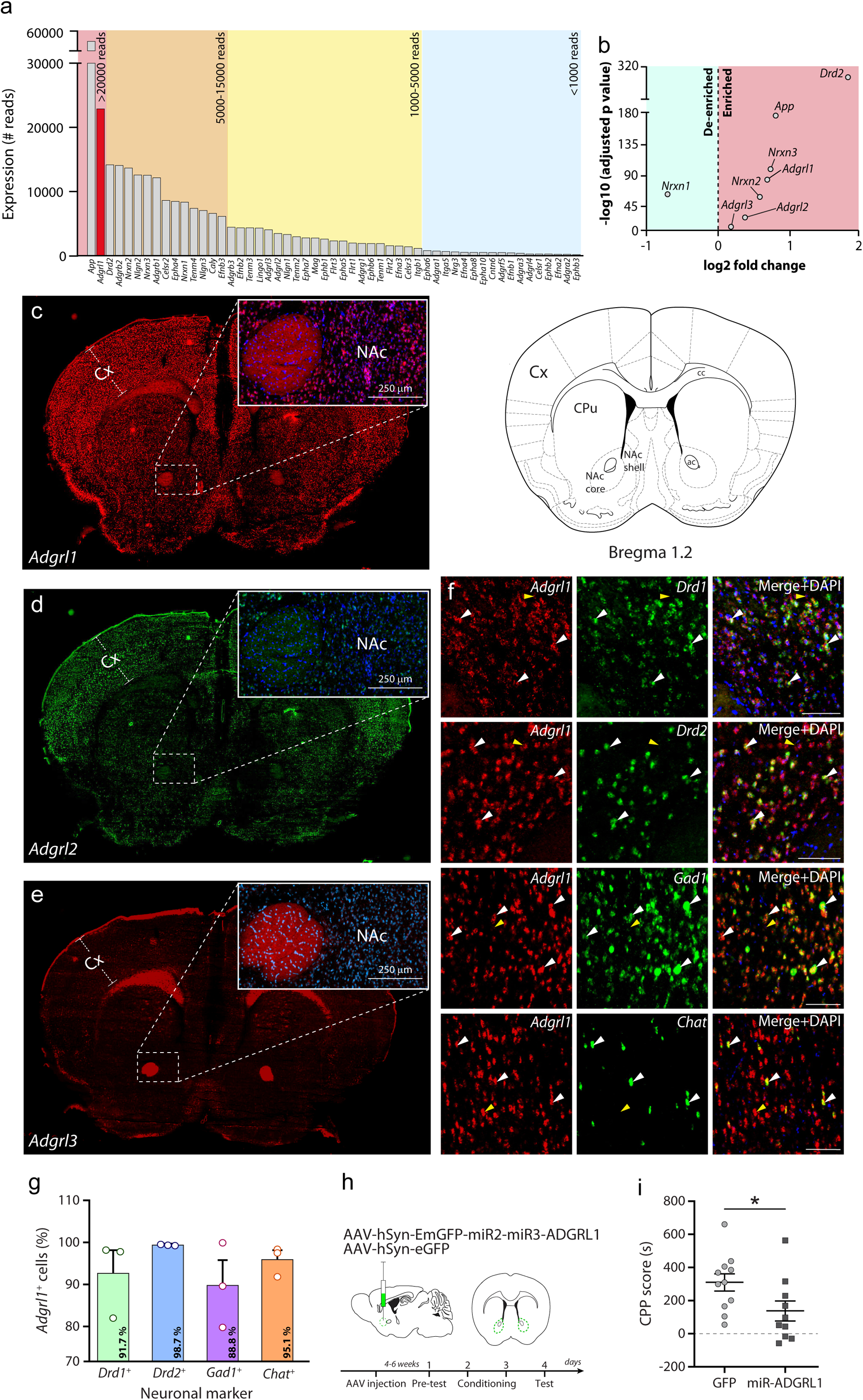
*Adgrl1* is an essential component of the nucleus accumbens’s cellular identity and function in cocaine reward. **a.** Actively translated mRNA coding for membrane proteins in D2 medium spiny neurons (MSNs), grouped from highest to lowest expression levels according to the number of RNAseq reads (left to right). Note that *Adgrl1* is the most expressed adhesion GPCR in D2-MSNs. **b.** Enrichment or de-enrichment of selected synaptic adhesion molecules in D2-MSNs. **c-e.** *Adgrl* isoforms mRNA expression in the mouse NAc using fluorescence in situ hybridization (FISH; *Adgrl1* in red, *Adgrl2* in green, *Adgrl3* in red) and DAPI for nuclear staining (in blue). Cx: cortex, cc: corpus callosum, CPu: caudoputamen, NAc core: nucleus accumbens core, NAc shell: nucleus accumbens shell, ac: anterior commissure. **f.** Identification of *Adgrl1* in specific neuronal subtypes in the NAc. Dopamine receptors type-1 and type-2: *Drd1* and *Drd2*, respectively, *Gad1*: glutamic acid decarboxylase 1, *Chat*: choline acetyltransferase. White arrowheads: positive cells for both markers; yellow arrowheads: *Adgrl1*-only positive cells **g.** Percentage of *Adgrl1*-positive cells in NAc neuronal subpopulations. >2400 cells per marker on 3 sections were analyzed (Qupath 0.3.2 software). **h.** Timeline of the CPP assay in C57BL6J mice injected in the NAc with adeno-associated virus (AVV) expressing EmGFP along with miRNAs against ADGRL1 or eGFP as control. **i.** Rewarding effect of cocaine measured as CPP score in control and NAc-specific ADGRL1 KD mice. Data are represented as mean ± S.E.M, n=10-11. Student’s t-test, \**p*<0.05.

We further compared the whole-brain distribution of *Adgrl1* expression to *Adgrl2* and *Adgrl3* using fluorescent *in situ* hybridization (FISH). All three latrophilins mRNA were detected in the brain at distinct fluorescent intensities. *Adgrl1* was detected throughout the brain in cortical and subcortical areas, in particular the NAc (Figure 1c-e and Supplementary Figure 1a-c). In contrast, *Adgrl2* and *Adgrl3* were weakly detected in the NAc, but strongly detected in the hippocampus (Figure 1c-e and Supplementary Figure 1a-c). This distribution profile was also observed in previous reports and in publicly available databases (Supplementary Table 3) ^35^ ^36^ ^37^ ^38^.

To precisely identify the neuronal subtypes expressing *Adgrl1* in the NAc, we performed dual-labeling FISH assays by co-labeling with probes targeting molecular markers tagging specific subpopulations. *Adgrl1* was detected in D1 and D2-MSNs, as well as in GABAergic and cholinergic interneurons (Figure 1f). Quantification of *Adgrl1* positive neurons showed its expression in 91.7% of *Drd1*-positive cells, 98.7% of *Drd2*-positive cells, 88.8 % of glutamic acid decarboxylase 1 (*Gad1*) positive cells and 95.1 % in choline acetyltransferase (*Chat*) positive cells (Figure 1g). In contrast to the RiboTag sequencing, the sensitivity of FISH allowed us to detect *Adgrl1* in multiple neuronal types in the NAc. The expression of *Adgrl1* in MSNs suggests a new function for this receptor in NAc-dependent behavioral responses, such as in drug addiction.

Next, in order to analyze the consequences of *Adgrl1* invalidation on the rewarding properties of the drug of abuse cocaine, we sought to evaluate the contribution of ADGRL1 expression in an experimental paradigm of drug-seeking behavior in mice, using the conditioned place preference (CPP) assay. In this assay, wild-type mice associate cocaine injection with environmental cues and express preference towards them, thus spending more time inside the chambers linked to cocaine injection. We thus generated mice bearing ADGRL1-deficiency in NAc neuronal populations by microinjecting AAV to deliver microRNA sequences targeting common exons in Adgrl1 mRNA splice variants (Chr 8 NC 000074.7: 84 664 979-84 668 576) under the control of the synapsin promoter and conducted CPP assays (Figure 1h-i, Supplementary Figure 1d-f). Mice displaying downregulation of ADGRL1 expression in NAc neurons performed poorly in the CPP assays with less time spent within compartments associated with cocaine than GFP-expressing control animals (Figure 1i). These results suggested that ADGRL1 is an essential component of the neuronal response allowing the NAc to attribute rewarding properties to cocaine.

### Cocaine effector hevin is a novel ligand for ADGRL1

Given the metabotropic nature of ADGRL1, we speculated on the identity of a potential cocaine-induced cue modulating ADGRL1 function in the NAc. Increasing evidence indicates that alterations in astrocyte-secreted factors are involved in neuropsychiatric disorders ^39^. Hevin is an astrocyte-secreted protein implicated in cocaine-dependent changes in glutamatergic signaling and structural plasticity ^28^. Similar to ADGRL1, knockdown of hevin in accumbal astrocytes, reduced cocaine-rewarding properties ^28^. We postulated that ADGRL1 mediates its effects on cocaine reward through hevin, thereby suggesting that hevin may function as a ligand for ADGRL1.

We first tested whether hevin physically interacts with ADGRL1 using complementary approaches. An “in-cell” capture assay was performed to immunodetect the retention of secreted HA-tagged hevin on whole cell extracts from cells transfected with ADGRL1 construct (Figure 2a-c). ADGRL1-expressing cells displayed heightened retention of hevin (Figure 2b-c), thus suggesting an interaction with this aGPCR. As expected, cells expressing the previously described receptor neurexin1-alpha (NRX1α) exhibited higher hevin retention compared to control cells transfected with an empty vector (Figure 2b-c). Hevin possesses various high-affinity calcium binding sites and its binding properties to NRX1α depend on the presence of calcium ^40^ (Supplementary Figure 3). To test the involvement of calcium on hevin-ADGRL1 interaction, the assay was performed with a calcium-chelating agent, EGTA. Chelation of calcium did not show significant changes in the retention of hevin despite a slight increase in HA-tagged hevin immunodetection (Figure 2d-e). Since the retention of hevin to ADGRL1-expressing cells remained unaffected by calcium absence or presence, we opted to proceed with the experiments using standard extracellular calcium concentrations.

**FIGURE 2.**
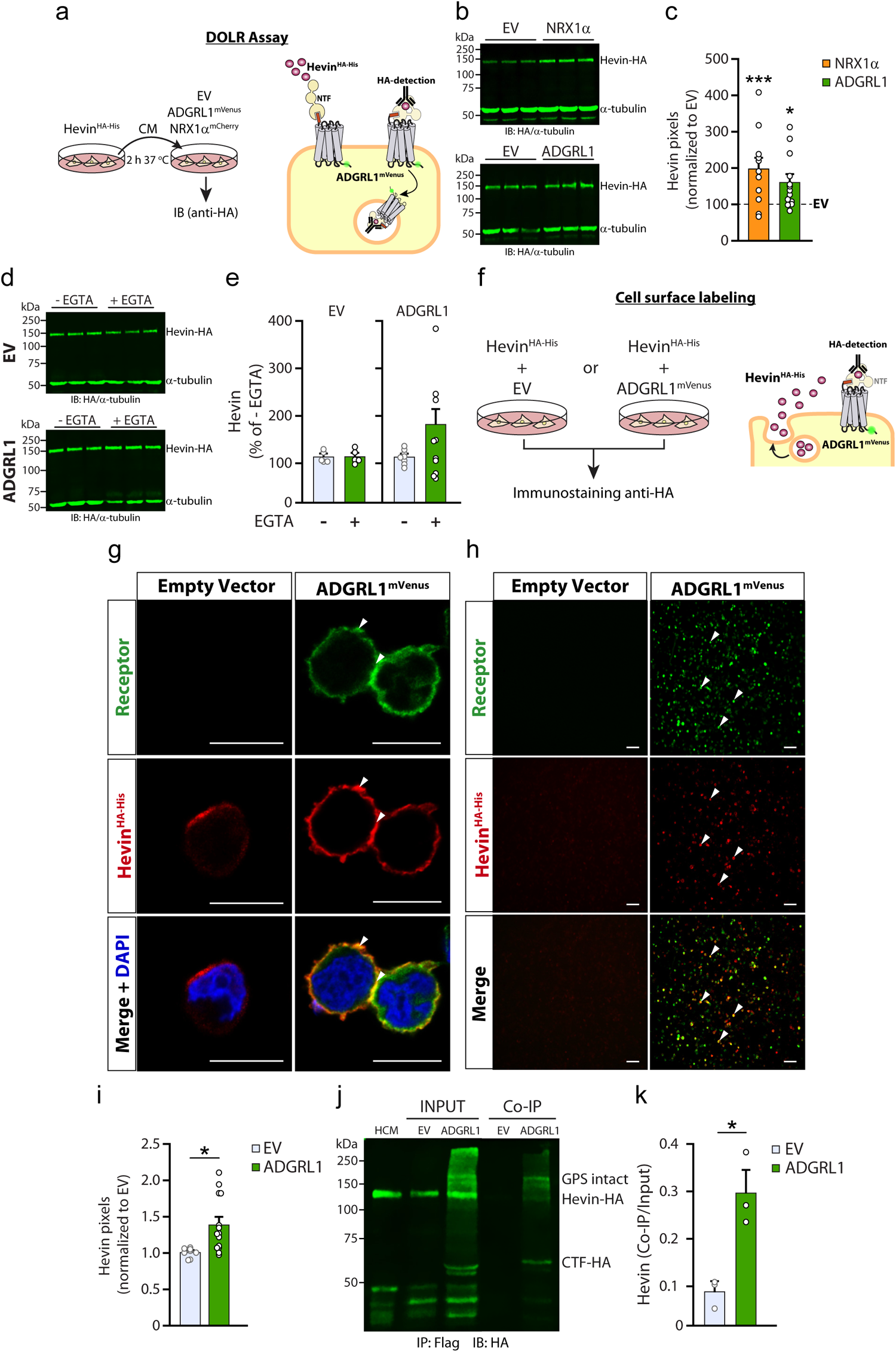
ADGRL1 binds to the cocaine effector hevin. **a.** Schematic representation of the DOLR assay. EV: empty vector, CM: conditioned media, IB: immunoblotting, NTF: N-terminal fragment. **b.** Representative fluorescent immunoblot of the DOLR assay, showing total extracts from HEK293T cells transfected with EV, NRX1α or ADGRL1 plasmids, after their incubation with hevin-containing conditioned media. Bands corresponding to HA-tagged hevin and α-tubulin as loading control are observed. Three lanes per sample were loaded. **c.** Quantification of fluorescence values for hevin immunodetection in b. **d.** Representative immunoblots from DOLR assays in the presence of EGTA. **e.** Quantification of fluorescence values for hevin immunodetection, expressed as a percentage of the condition without EGTA. **f.** Schematic representation of the cell-surface labeling assay. **g.** Representative confocal images from the cell-surface labeling assay to detect membrane-bound hevin^HA-His^ (HA tag, in red) onto cells co-transfected with EV or co-expressing ADGRL1^mVenus^ (in green). Merge panel also displays nuclear DAPI staining. Scale bar 10 µm. **h.** Epifluorescence microscopy of the cell-surface labeling assay seen in g. **i.** Quantification of hevin total pixels (red channel) from the images obtained in h. **j.** Representative immunoblot from co-immunoprecipitation assays. Whole-cell lysates from cells expressing ADGRL1^Flag-HA^ or mock-transfected exposed to hevin-containing media were incubated with beads coupled to anti-Flag antibody. HA-tagged hevin co-immunoprecipitation with Flag-tagged ADGRL1 was detected by anti-HA immunodetection. **k.** Input-normalized fluorescence values of hevin-immunodetected bands obtained in i. Data in c,e,i,k are represented as mean ± S.E.M of at least three independent experiments. Student’s t-test, \**p*<0.05, \*\*\**p*<0.001.

The binding of hevin to the cell surface of ADGRL1-expressing cells was further assessed by confocal microscopy using a cell-surface labeling assay. mVenus-tagged ADGRL1 and HA-tagged hevin were co-transfected and immunolabeled with anti-HA and fluorescent secondary antibodies in non-permeabilizing conditions (Figure 2f). Initially, we struggled to detect any signal for hevin when fixing conditions involved the absence of CaCl_2_ and MgCl_2_, in phosphate saline buffer, hence technical changes were made to achieve hevin detection (methods section). A faint fluorescent signal for hevin was detected on the surface of EV-transfected cells but was significantly higher when monitored for cells expressing the positive control NRX1α (Figure 2g, Supplementary Figure 4). Similarly, ADGRL1-expressing cells displayed a higher fluorescent signal corresponding to hevin than control cells, a result consistent with receptor-ligand interaction (Figure 2h-i). Protein-protein complex formation between ADGRL1 and hevin was also monitored by conducting a co-immunoprecipitation assay. Cells expressing Flag-tagged ADGRL1 were incubated with hevin-containing media and subsequently lysed to affinity-purify ADGRL1 using beads coupled to an anti-Flag antibody. Immunoprecipitation of ADGRL1 resulted in a concomitant enrichment of hevin (Figure 2j-k), confirming their association within the same protein complex and indicating that ADGRL1 serves as a receptor molecule for hevin.

### Hevin modulates ADGRL1 adhesion properties through receptor stability/assembly and internalization

As a synaptic adhesion molecule, ADGRL1 contributes to the stabilization of synapses through the function of its extracellular adhesion motifs ^41^. Hence, we evaluated whether hevin could affect intercellular adhesion junctions elicited by ADGRL1 and its cognate ligands. We conducted cell aggregation assays which allow to reveal cell-cell interactions through complementary binding of membrane-embedded adhesion molecules in trans. Individual cell populations were mixed, each expressing either ADGRL1 or its ligand NRX1β, along with fluorescent markers GFP or DsRed respectively, and the formation of cell aggregates was monitored in the presence or absence of hevin in the media (Figure 3a). While ADGRL1/NRX1β-dependent cell aggregates display a size increase in the absence of hevin during the time course of the experiment, the presence of hevin limits the magnitude of growth and frequency of high-order aggregates, denoted by a lower aggregation index (Figure 3a-d, Supplementary figure 5). These data indicate that hevin exerts a destabilization effect on adhesion complexes mediated by ADGRL1 (Figure 3b-d).

**FIGURE 3.**
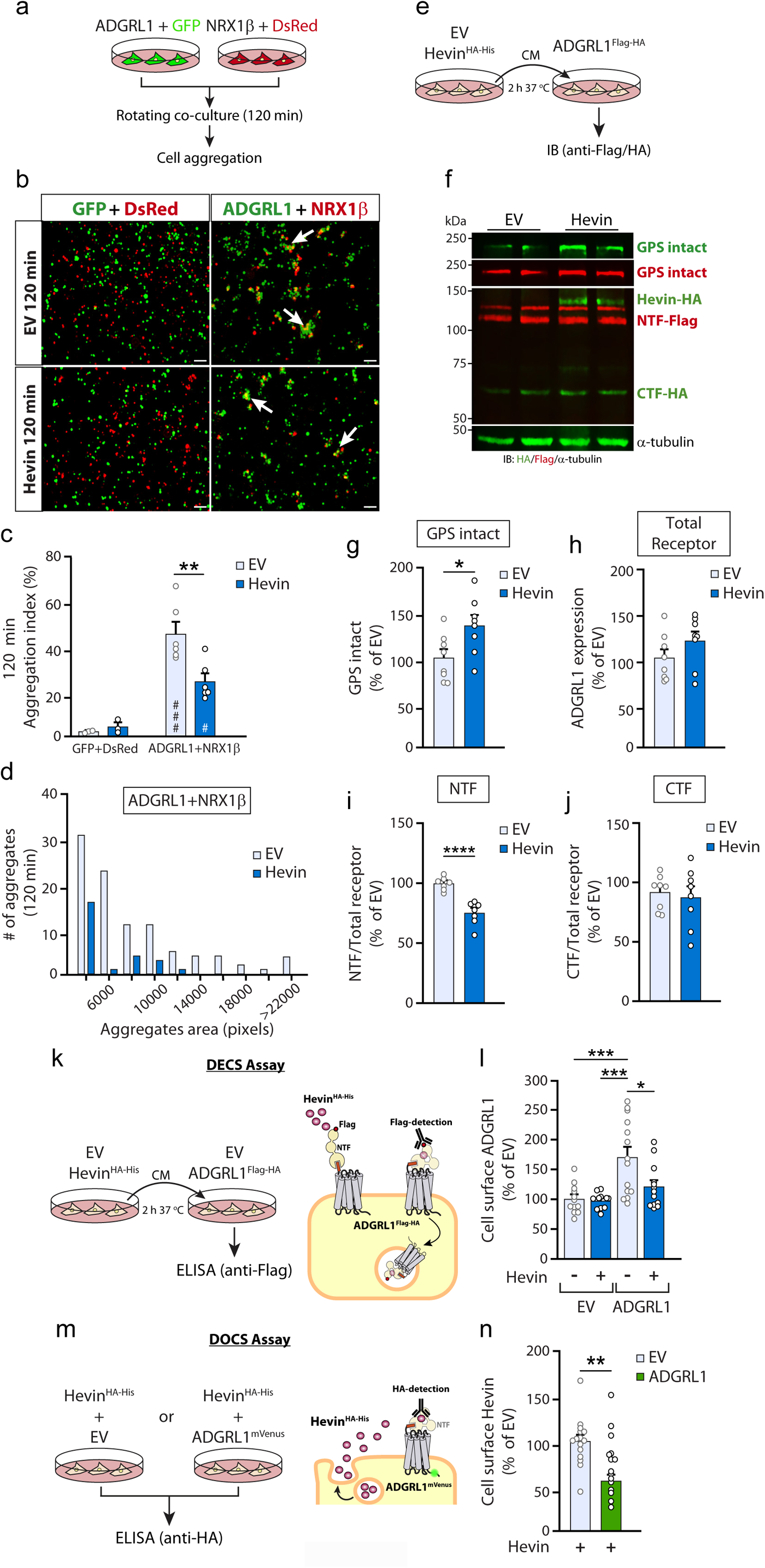
Hevin modifies the adhesive properties of ADGRL1 through modulation of receptor fragments stability and internalization. **a.** Schematic representation of cell aggregation assays. Cells co-expressing ADGRL1 and GFP or pCMV-NRX1β and DsRed, were mixed and incubated in the absence or presence of hevin-containing media. **b.** Representative images from cell aggregation assays. **c.** quantification of aggregation index of cell aggregation assays conducted in b. **d.** Size distribution of aggregates obtained in b. **e.** Schematic representation of DOLR assay for the detection of ADGRL1. CM: conditioned media, IB: immunoblotting. **f.** Representative immunoblot for the assay represented in e. ADGRL1-expressing cells were exposed to hevin-containing media or control media and whole-cell lysates were probed with anti-HA antibody to detect HA-tagged Hevin and ADGRL1-CTF, or anti-Flag antibody to detect Flag-tagged ADGRL1-NTF. The fraction of ADGRL1 not processed through the GPS (GPS intact) was detected using both anti-HA and anti-Flag. α-tubulin was detected as a loading control. Two lanes per sample were loaded. NTF/CTF: N-/C-terminal fragments respectively. **g-j.** Normalized fluorescence values for bands representing GPS-intact ADGRL1 (g), ADGRL1 total expression levels (h) NTF proportion (i) and CTF proportion (j). **k.** Schematic representation of the DECS assay detecting surface-exposed receptor with HRP-coupled secondary antibody against anti-Flag in ADGRL1-expressing cells exposed to hevin-containing media. EV: empty vector, CM: conditioned media. **l.** Quantification of absorbance values resulting from DECS assays normalized to EV condition. **m.** Schematic representation of the DOCS assay. **n.** Normalized hevin levels detected on the surface of cells co-expressing ADGRL1^mVenus^ or mock-transfected cells. Data are represented as mean ± S.E.M of at least three independent experiments. One-way ANOVA and Dunnet test (l), Student’s T test (c, g-j,n), \**p*<0.05, \*\**p*<0.01, \*\*\**p*<0.001. #: significant difference compared to GFP+DsRed condition (c).

The hallmark autoproteolysis event occurring at a consensus site named GPS in many aGPCRs, has been reported to modulate their function ^42^ ^43^ ^44^. Hence, we evaluated the impact of hevin on receptor cleavage into its two receptor adjuncts indicated as an N-terminal fragment (NTF) and a C-terminal fragment (CTF). ADGRL1-expressing cells were incubated in the presence or absence of hevin and subsequently analyzed by immunoblotting using antibodies against epitope tags positioned on the two separate fragments (Figure 3e-f). While total receptor levels displayed a slight increase when cells were exposed to hevin, this was not significant (Figure 3h). Furthermore, total NTF and CTF levels remained unchanged in both conditions (Supplementary Figure 6c,d). Nonetheless, the levels of NTF in relation to total receptor levels were decreased suggesting a destabilizing effect on the half-life of this receptor fraction in contrast to CTF levels which did not experience alterations (Figure 3i-j). This immunoblotting strategy allowed us to detect an additional receptor fraction representing entities that were not cleaved at GPS and thus in which it was kept intact. Surprisingly, immunodetection of this uncleaved receptor fraction was increased, suggesting that its expression was stabilized upon hevin exposure (Figure 3g).

Sustained ligand exposure often leads to GPCR internalization and degradation, a process resulting from functional ligand-receptor interactions ^45^. Thus, given the destabilizing nature of hevin on ADGRL1 fragments, we sought to determine whether the complex was internalized upon sustained exposure. Cells expressing ADGRL1 and incubated in hevin-containing media for 2 hours were monitored for receptor cell-surface expression using immunodetection. Prolonged hevin exposure reduced the levels of ADGRL1 at the cell surface as compared to control conditions in absence of hevin (Figure 3k-l). Concomitantly, detection of cell-surface levels of hevin was decreased in presence of ADGRL1 (Figure 3m-n). The latter was accompanied by a higher level of hevin detection in whole-cell extracts contrasting with lower secretion levels of hevin when ADGRL1 was present, results that are consistent with increased intracellular hevin accumulation mediated by ADGRL1 (Figure 2b-c, Supplementary Figure 7).

### Hevin drives ADGRL1-G protein signaling bias

Given that GPCR internalization results from sustained receptor activation, we sought to elucidate the G protein activation pattern of ADGRL1 when exposed to the internalization-inducing agent hevin. For this, we employed the recently developed TRUPATH BRET-based biosensors as a tool for evaluating the coupling of various G proteins associated with major metabotropic pathways ^30^. These biosensors consist of the Gα-Gβ-Gγ protein trimer, where Gα is coupled to Rluc8 and Gγ to GFP2. Under conditions where the G protein is inactive, the trimer is stabilized and GFP2 fluorescence emission can be elicited by excitation through the light-producing Rluc8 due to their proximity, thus resulting in a high BRET^2^ signal. Conversely, the activated G protein biosensor yields a decrease in BRET^2^ signal due to the dissociation of Gα (Figure 4a). Therefore, we initially characterized the G protein coupling profile of ADGRL1 linked to its constitutive activity in living cells. HEK293 cells were co-transfected with the BRET-based G protein biosensors and increasing amounts of the receptor plasmid to obtain a gradation of expression levels, along with empty vector to maintain a constant DNA load in all conditions. The high BRET^2^ signal levels that was detected for basal conditions in the absence of receptor expression were gradually lowered when increasing receptor plasmid concentrations were used, and this for all G protein biosensors tested being Gi, Gs, G12/13 and Gq/11 (Figure 4b-g, Supplementary Figure 8). The BRET^2^ signal dynamic range differed slightly between biosensors with the most pronounced effect detected for Gs and G12. However, the biosensors’ activation kinetics induced by ADGRL1 revealed a more sensitive coupling to Gs giving that BRET^2^ saturation levels were reached using lower receptor plasmid concentrations (50ng) as compared to other biosensors requiring higher plasmid concentrations (250ng or more).

**FIGURE 4.**
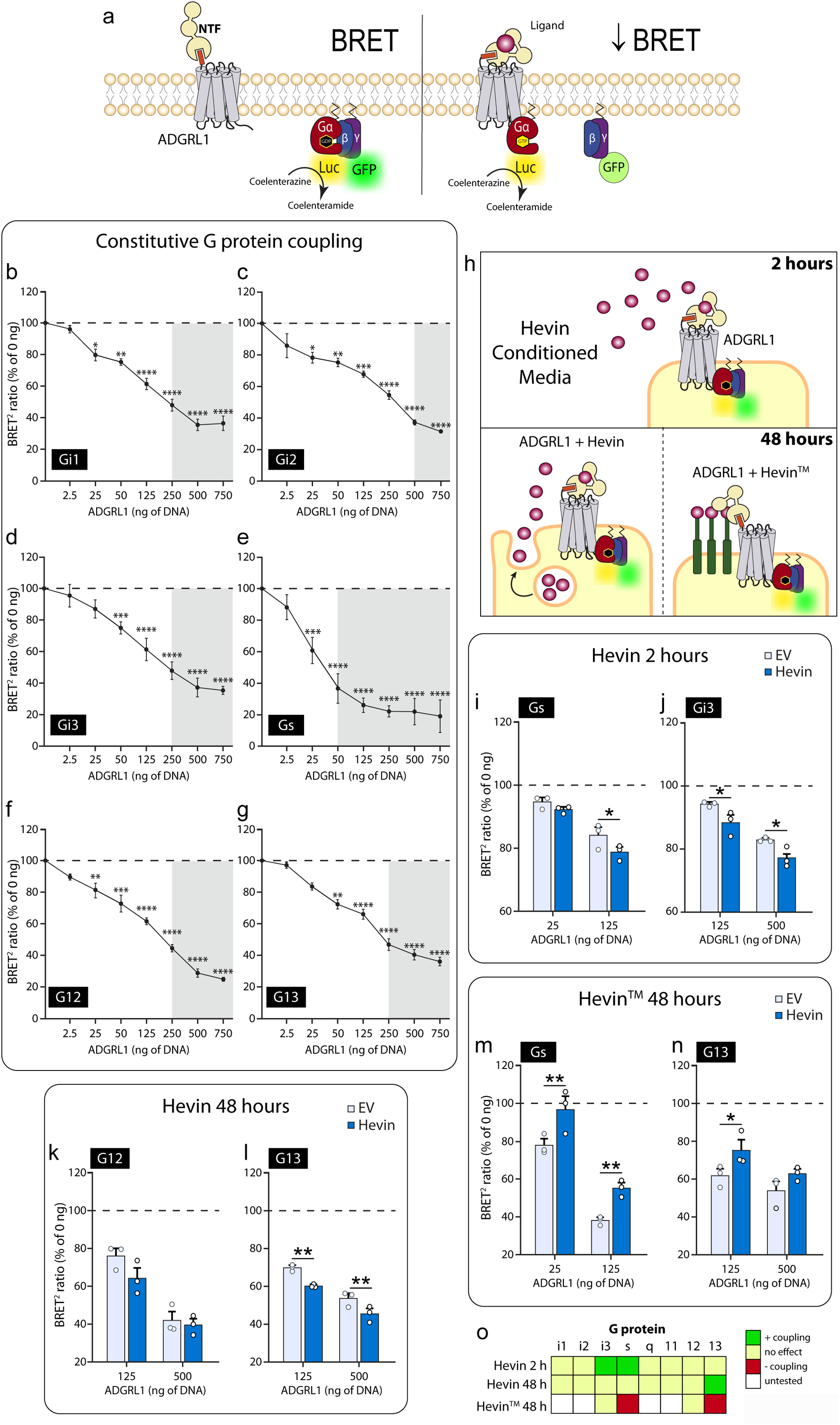
ADGRL1 signaling is biased toward selected G proteins upon exposure to hevin. **a.** Schematic representation of BRET-based G protein biosensors used to evaluate the coupling of ADGRL1 to G proteins. **b-g.** Constitutive coupling profile of ADGRL1 to Gi1, Gi2, Gi3, Gs, G12 and G13. **h.** Schematic representation of the experimental settings used to assess the effect of hevin on ADGRL1 G-protein coupling profile, monitored using BRET-based G protein biosensors. In the top panel, cells co-expressing ADGRL1^Flag-HA^ and biosensors were incubated for 2 h with conditioned media from Hevin^HA-His^ -expressing cells, before BRET^2^ measurements were initiated. In the bottom left and right panels, BRET^2^ measurements were conducted using cells co-expressing ADGRL1^Flag-HA^, G protein biosensors, and either Hevin^HA-His^ or Hevin^TM^. **i-j.** Normalized BRET^2^ ratio values of the assay represented in h (top) for Gs and Gi3 biosensors. **k-l.** Normalized BRET^2^ ratio values of the assay represented in h (bottom left) for G12 and G13 biosensors. **m-n.** Normalized BRET^2^ ratio values of the assay represented in h (bottom right) for Gs and G13 biosensors. **o.** Heatmap displaying the differential effect of hevin exposure on ADGRL1 G-protein coupling profile. Data are represented as mean ± S.E.M of at least three independent experiments. Two-way ANOVA, \**p*<0.05, \*\**p*<0.01 \*\*\**p*<0.001, \*\*\*\**p*<0.0001.

We then assessed whether hevin could modify the G protein coupling profile of ADGRL1. Cells co-expressing ADGRL1 and G protein biosensors or biosensors only were incubated for 2 h with hevin or control media, followed by BRET^2^ measurements (Figure 4h top). Hevin selectively promoted the coupling of ADGRL1 to Gs and Gi3, which was reflected by a more pronounced decrease in the BRET^2^ ratio compared to basal activity (cells treated with control media) (Figure 4i-j). No changes in BRET^2^ signals were detected for the remaining G proteins tested whether ADGRL1 was expressed or not (Figure 4o, Supplementary Figures 9,10a). These results indicate that hevin-ADGRL1 signaling possesses a bias component.

Because the extent of ligand exposure profoundly influences GPCR coupling to G proteins ^46^ ^45^, we also examined the effect of chronic hevin exposure on ADGRL1 signaling. Cells co-expressing ADGRL1 and hevin along with G protein biosensors were kept in culture for a 2 day-period and monitored using BRET^2^ measurements (Figure 4h, bottom left). The presence of hevin solely affected ADGRL1 activation of G13 biosensor by favoring the magnitude of coupling (Figure 4l), without significant changes observed for the same family member G12 (Figure 4k), the acutely activated Gs and Gi3, nor the remainder of G protein biosensors evaluated (Figure 4i, Supplementary Figure 10b). Using another paradigm of hevin long-term exposure, we increased its bioavailability into membrane compartments therefore constraining it to all cellular trafficking routes taken by ADGRL1. We thus co-transfected cells with a construct encoding hevin fused to a synthetic transmembrane domain along with the plasmids encoding ADGRL1 and G protein biosensors, and conducted BRET^2^ measurements following a 2-day post-transfection period (Figure 4h, bottom right, Supplementary Figure 10c). Interestingly, this configuration caused an inhibitory effect on the coupling of ADGRL1 to acutely-activated Gs and to chronically-activated G13, suggesting that not only the extent but also the compartmentalization of hevin exposure can fine-tune the bias of ADGRL1 signaling profile.

## DISCUSSION

Alterations in the nucleus accumbens constitute the foundation for aberrant neuronal connectivity in cocaine addiction, where the natural reward response is hijacked. Here we describe the morphology-sensitizing GPCR ADGRL1/latrophilin-1 as a new modulator for cocaine reward. We demonstrate that this receptor acts as a sensor for the astrocytic protein hevin, which modulates ADGRL1 adhesion properties by affecting its stability, internalization and signaling.

Knocking down ADGRL1 specifically in adult NAc neurons caused a decrease in the rewarding properties of cocaine, demonstrating that ADGRL1 expression in NAc is required for the rewarding effects of this drug. The nucleus accumbens receives dopaminergic, GABAergic and glutamatergic inputs from other brain regions ^47^. Since ADGRL1 is essential for proper excitatory and inhibitory synapse function ^10^ ^48^, it could be modulating cocaine-evoked synapses within the NAc in specific neuronal subtypes. Our analysis of neuron-specific mRNA sequencing in the NAc shows that ADGRL1 is strongly enriched in D2 MSNs compared to all other neuronal subtypes, suggesting that ADGRL1’s effects on cocaine reward could be generated via its effect on D2 MSNs ^34^. However, the receptor is also expressed in other neuronal populations of NAc, including D1 MSNs and interneurons which are also implicated in the regulation of the response to cocaine ^49^ ^50^. Hence, we cannot exclude an effect of ADGRL1 on cocaine-rewarding properties via other neuronal subtypes in NAc. The targeting of ADGRL1 knockdown to specific neuronal subpopulations in the adult NAc would help dissect how distinct cellular context impacts receptor contribution. Furthermore, while we cannot predict the role of ADGRL1 in motivation to consume substances of abuse, interestingly, deficiency of ADGRL1 in mice results in obesity originating from increased food consumption, suggesting a role for ADGRL1 in motivation to control food intake ^51^. Importantly, ADGRL1 might be essential for behavior directed toward natural reward such as social behavior which is impaired in individuals with ASD and that seems to arise from an aberrant reward response ^52^ ^53^.

We explored which cocaine-induced signal could be modulating ADGRL1 function. Neurotransmission is strongly influenced by astrocytic secreted factors ^54^ ^55^ ^56^ because of the close proximity of astrocytic processes to nearly every glutamatergic synapse ^57^. Recent studies demonstrate that alterations in glycoproteins secreted from astrocytes are implicated in rodent models of neuropsychiatric disorders including cocaine addiction ^9^ ^28^. While most astrocytic factors are expressed only during development, hevin remains highly expressed in the adult ^58^, where it has been shown to be essential for cocaine drug-seeking behavior. The role of hevin on synaptic connectivity was previously attributed to its interaction with neurexins and neuroligins, however, neurexins and neuroligins knockouts did not prevent hevin from regulating synapse function ^59^, suggesting that another receptor might be mediating the function of hevin on cocaine reward.

Using complementary approaches, we found that hevin is a ligand for ADGRL1 and that it stabilizes the receptor’s full-length form while destabilizing the receptor’s adhesion motif-bearing extracellular NTF, which might directly affect its function. ADGRL1 ectodomain contains an olfactomedin domain. Hevin C-terminal has been reported to bind the olfactomedin domain of the protein myocilin ^60^ ^61^, suggesting that it could bind to ADGRL1 via a similar protein interaction. The relevance of the hevin-ADGRL1 interaction is also highlighted by the disruptive effect of hevin on the adhesion of ADGRL1 to its previously characterized ligand NRX1β, the binding of which occurs through the olfactomedin domain. The binding of hevin also induced ADGRL1 internalization, and hevin was simultaneously internalized with it. Surface receptor internalization is an essential process in cell physiology for the regulation of GPCR-mediated signaling ^62^. While classical concepts view receptor internalization as a mechanism to end GPCR-mediated signaling, recent studies show that GPCRs can signal from endosomes by forming a complex with specific G proteins and β-arrestins ^63 64 65 66^.

ADGRL1 has previously been shown to couple to Gi, Gs, Gq and G12/13 G proteins ^13^ ^11^ ^67^ ^68^. Here, using the novel BRET biosensors technology in living cells, we validated that ADGRL1 can constitutively couple to different G proteins. This receptor was reported to exhibit basal activity ^67^, which can be modulated upon ligand exposure ^13^. Short-term exposure to hevin enhances ADGRL1’s activation of Gi3 and Gs, whereas prolonged exposure promotes its coupling to G13. The differential coupling to various G proteins at these distinct time points hints at a biphasic response mode in hevin’s modulation of ADGRL1 activity. Activation of Gs and G13 have been shown to regulate mainly the levels of cAMP and Rho GTPases, respectively ^69^. The signaling pathways of these G proteins are required for the activation of the transcriptional factors CREB and SRF, both capable of inducing the expression of ΔFosB ^70^, a gene responsible for many of the long-term cocaine-induced changes in synaptic plasticity and behavior ^71^. Meanwhile, the second phase involving G13-dependent signaling on actin cytoskeleton would reinforce and/or increase the structural changes and connections formed in the first phase through Gs signaling. These results support the hypothesis that hevin regulates drug-related processes via its binding to ADGRL1 (Figure 5). However, we did not explore hevin-ADGRL1 interaction directly in the NAc, due to complexities intrinsic to the physiological context, such as ligand competition for either hevin or ADGRL1, as well as the possible occurrence of compensatory mechanisms in response to genetic manipulations. Further studies will be required to address this point.

**FIGURE 5.**
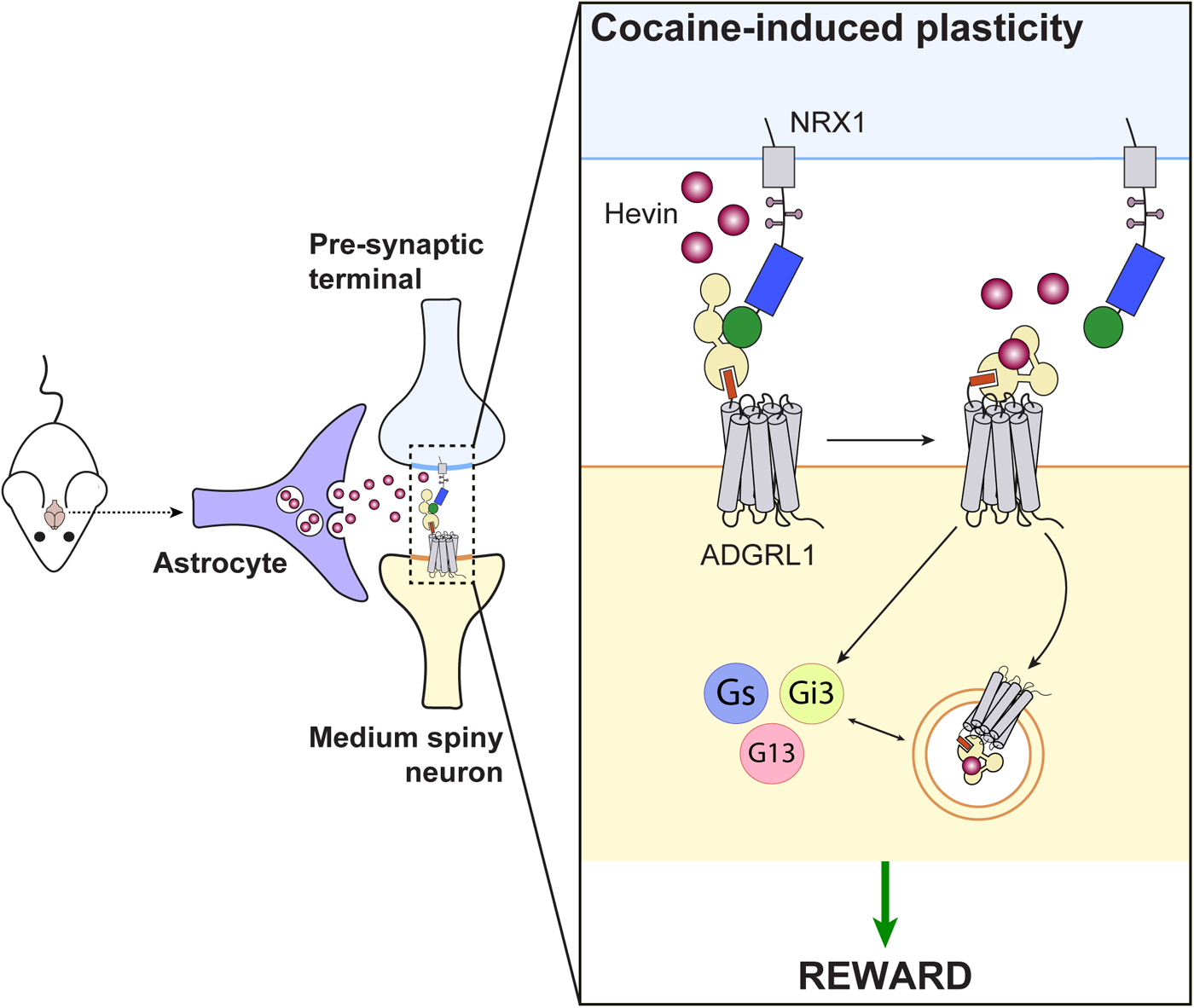
ADGRL1 senses astrocytic cues associated with cocaine reward. A model proposing how hevin sensing by ADGRL1 could modulate cocaine reward. Cocaine consumption may result in MSN-expressed ADGRL1 binding the astrocytic protein hevin subsequently inducing intracellular signaling through the modulation of Gs/Gi/G13 proteins activity. Moreover, hevin also affects intercellular adhesion complexes between ADGRL1 and neurexin thus supporting neuronal plasticity.

To conclude, we show for the first time a role for ADGRL1, an adhesion GPCR, in substance abuse disorder. We also describe a novel interaction for both ADGRL1 and hevin, which suggests a possible new communication axis between astrocytes and neurons in the context of reward-related behaviors. Future studies will further explore the implications of this novel ligand-receptor pair in terms of neuronal plasticity, with an emphasis on structural, transcriptional and connectivity changes, not only in the NAc, but also in other brain regions related to the reward circuit. Since ADGRL1 is associated with several neuropsychiatric disorders, our work highlights the relevance of this family of proteins as a susceptibility hub underlying reward-related comorbidity.

## Acknowledgments

We thank Juana Amador Calderon, Ricardo A. Bautista Torres and Franck Louis for technical assistance. We extend our gratitude to Cagla Eroglu (Duke University School of Medicine) for providing hevin constructs. We are also grateful to Monserrat Avila-Zozaya and Ana L. Moreno-Salinas for guidance in confocal microscopy and BRET assays, respectively. This work was supported by grants to AAB from Consejo Nacional de Humanidades, Ciencia y Tecnología (CONAHCYT; #CB221568, #PN2017-4687, Infra #302758), Secretaría de Educación Pública (SEP)-Cinvestav (#233), Secretaría de Educación, Ciencia, Tecnología e Innovación (SECTEI; #SECTEI/165/2023) and Munck-Pfefferkorn mentoring award, as well as a joint CONAHCYT-ECOS Nord grant to AAB and VV (SEP-CONAHCYT-ANUIES-ECOS Nord #296652 [MEX], #M18S02 [FRA]). The study received additional funding to VV from Institut National de la Santé et de la Recherche Médicale (INSERM), Centre National de la Recherche Scientifique (CNRS), Sorbonne Université, the Brain & Behavior Research Foundation (NARSAD Young Investigator Award, #17566), FP7 Marie Curie Actions Career Integration Grant (FP7-PEOPLE-2013-CIG 618807), Promouvoir l’Excellence de la Recherche à Sorbonne Université (PER-SU 2014), Agence Nationale de la Recherche (ANR JCJC 2015 ANR-15-CE16-0005, Hevinsynapse). This article was part of the PhD thesis of KGCB (CONAHCYT doctoral scholarship #723831).

## Authors contribution

Research was performed by KGCB, AA and SD; data mining was conducted by EV; data were analyzed by KGCB, AA, EV, NH, VV and AAB; research was supervised by VV and AAB; manuscript was written, revised and corrected by KGCB, SD, EV, NH, VV and AAB; figures were elaborated by KGCB, AA, VV and AAB.

## Conflict of Interest

Authors declare no conflict of interest.

## Data Availability

Products of this article can be made available from the corresponding author upon reasonable request.

## Notes

### Competing Interest Statement

The authors have declared no competing interest.

