## Supplementary information for "Genetic liability underlying reward-related comorbidity in psychiatric disorders involves the coincident functions of autism-linked ADGRL1 and hevin"

**Plasmids**

pCMV-Adgrl1^Flag-HA^, encoding rat Adgrl1 with the Flag tag in its amino-terminal fragment and HA in the first extracellular loop, was previously described ^1^. Adgrl1 used in this study corresponds to the following splicing variant: XM_039097967.1.

pCMV-Adgrl1^mVenus^ encoding rat Adgrl1 coupled to mVenus at its carboxyl-terminal fragment was previously described ^2^.

pCMV-NRX1α^Flag-mCherry^ encoding for the rat Nrx1α without the SSA splicing site, with the Flag tag in its amino-terminal region and coupled to mCherry in its carboxyl-terminal region ^3^.

pCMV-NRX1β^Flag^ encoding for the rat NRX1β, with the Flag tag in its amino-terminal region ^4^.

pCMV-Hevin^HA-His^ encoding mouse hevin fused with HA and poly Histidine tags at its carboxyl-terminal (Genecopoeia, Ex-Mm-34208-08).

pCMV-Hevin^HA-TM^ encoding mouse hevin fused with HA and the transmembrane region of LDL and the cytoplasmic tail of CD46 was previously described ^5^.

pcDNA™6.2-GW/EmGFP-miR1, pcDNA™6.2-GW/EmGFP-miR2 and pcDNA™6.2-GW/EmGFP-miR3 encoding iRNA against Adgrl1 were obtained by cloning antisense sequences described in Supplementary Table 1 into the pcDNA™6.2-GW/EmGFP vector, according to the manufacturer’s instruction (BLOCK-iT™ Pol II miR RNAi Expression Vector Kit with EmGFP).

pAAV-hSyn-eGFP encoding for the enhanced green fluorescent protein (eGFP) under the human synapsin promoter (Addgene, 50465).

**Supplementary Table 1. Oligos for the generation of oligonucleotide probes.**

| **Gene** | **Oligo** | **Sequence (T3/T7 promoter)** |
| --- | --- | --- |
| *Adgrl1* | Forward | AATTAACCCTCACTAAAGGGACTATGCCAGCGGGGCCAACC |
|  | Reverse | TAATACGACTCACTATAGGGCTCAGGGAGCCCCAGACCCC |
| *Adgrl2* | Forward | AATTAACCCTCACTAAAGGGAGGCGTGCAGGTCGTGGACTG |
|  | Reverse | TAATACGACTCACTATAGGGAACATGGCCGGGAGGGAGGG |
| *Adgrl3* | Forward | AATTAACCCTCACTAAAGGGACACGAGCGCTCCAGCGAACA |
|  | Reverse | TAATACGACTCACTATAGGGTGGCTGGGCAGCACTCCAAT |

**Supplementary Table 2. ssOligos for miRNAs generation.**

| **miRNA** | **Oligo** | **Sequence** |
| --- | --- | --- |
| miR1 | Top | TGCTGGCAATTCCTCGAGTTCCCTTGGTTTTGGCCACTGACTGACCAAGGGAACGAGGAATTGC |
|  | Bottom | CCTGGCAATTCCTCGTTCCCTTGGTCAGTCAGTGGCCAAAACCAAGGGAACTCGAGGAATTGCC |
| miR2 | Top | TGCTGTGTTGTGCACCAGCTCTGAGAGTTTTGGCCACTGACTGACTCTCAGAGGGTGCACAACA |
|  | Bottom | CCTGTGTTGTGCACCCTCTGAGAGTCAGTCAGTGGCCAAAACTCTCAGAGCTGGTGCACAACAC |
| miR3 | Top | TGCTGAGTGACCAACTGCATTTGCCCGTTTTGGCCACTGACTGACGGGCAAATAGTTGGTCACT |
|  | Bottom | CCTGAGTGACCAACTATTTGCCCGTCAGTCAGTGGCCAAAACGGGCAAATGCAGTTGGTCACTC |

**Supplementary Table 3. Relative expression between latrophilins according to different databases**

| **Database** | **Region** | **Adgrl1** | **Adgrl2** | **Adgrl3** |
| --- | --- | --- | --- | --- |
| Allen Brain Atlas | STR | + | ++ | +++ |
| Dropviz^m^ | STR (D1+) | +++ | ++ | ++ |
| Dropviz^m^ | STR (D2+) | +++ | + | ++ |
| Expression Atlas^m^ | STR | +++ | ++ | + |
| Expression Atlas^h^ | NAc | +++ | + | ++ |
| GTEx Portal^h^ | NAc | +++ | + | ++ |

STR: striatum, NAc: nucleus accumbens, m: mouse data, h: human data. The data used for the analyses described in this table were obtained from the GTEx Portal on 10/04/23.

**Supplementary Table 4. List of genes shown in Figure 1a-b.**

| **Gene symbol** | **Common name** | **NCBI Gene ID** |
| --- | --- | --- |
| *App* | Amyloid beta recursor protein | 11820 |
| *Adgrl1* | Latrophilin 1 | 330814 |
| *Drd2* | Dopamine receptor D2 | 13489 |
| *Adgrb2* | Brain-specific angiogenesis Inhibitor (Bai) 2 | 230775 |
| *Nrxn2* | Neurexin 2 | 18190 |
| *Nlgn2* | Neuroligin 2 | 216856 |
| *Nrxn3* | Neurexin 3 | 18191 |
| *Adgrb1* | Brain-specific angiogenesis Inhibitor (Bai) 1 | 107831 |
| *Celsr2* | Cadherin, EGF LAG seven-pass G-type receptor 2 | 53883 |
| *Epha4* | Eph receptor A4 | 13838 |
| *Nrxn1* | Neurexin 1 | 18189 |
| *Tenm4* | Teneurin transmembrane protein 4 | 23966 |
| *Nlgn3* | Neuroligin 3 | 245537 |
| *Caly* | Calcyon | 68566 |
| *Efnb3* | Ephrin B3 | 13643 |
| *Adgrb3* | Brain-specific angiogenesis Inhibitor (Bai) 3 | 210933 |
| *Efnb2* | Ephrin B2 | 13642 |
| *Tenm3* | Teneurin transmembrane protein 3 | 23965 |
| *Lingo1* | LINGO-1 | 235402 |
| *Adgrl3* | Latrophilin 3 | 319387 |
| *Adgrl2* | Latrophilin 2 | 99633 |
| *Nlgn1* | Neuroligin 1 | 192167 |
| *Tenm2* | Teneurin transmembrane protein 2 | 23964 |
| *Epha7* | Eph receptor A7 | 13841 |
| *Mag* | Myelin-associated glycoprotein | 17136 |
| *Ephb1* | Eph receptor B1 | 270190 |
| *Flrt3* | Fibronectin leucine-rich transmembrane protein 3 | 71436 |
| *Epha5* | Eph receptor A5 | 13839 |
| *Flrt1* | Fibronectin leucine-rich transmembrane protein 1 | 396184 |
| *Adgrg1* | Adhesion G protein-coupled receptor G1 | 14766 |
| *Ephb6* | Eph receptor B6 | 13848 |
| *Tenm1* | Teneurin transmembrane protein 1 | 23963 |
| *Flrt2* | Fibronectin leucine-rich transmembrane protein 2 | 399558 |
| *Efna3* | Ephrin A3 | 13638 |
| *Celsr3* | Cadherin, EGF LAG seven-pass G-type receptor 3 | 107934 |
| *Itgb1* | Integrin beta 1 | 16412 |
| *Epha6* | Eph receptor A6 | 13840 |
| *Adgra1* | Adhesion G protein-coupled receptor A1 | 52389 |
| *Itga5* | Integrin alpha 5 | 16402 |
| *Nrg3* | Neuregulin 3 | 18183 |
| *Efna4* | Ephrin A4 | 13639 |
| *Epha8* | Eph receptor A8 | 13842 |
| *Epha10* | Eph receptor A10 | 230735 |
| *Cntn6* | Contactin 6 | 53870 |
| *Adgrf5* | Adhesion G protein-coupled receptor F5 | 224792 |
| *Efnb1* | Ephrin B1 | 13641 |
| *Adgra3* | Adhesion G protein-coupled receptor A3 | 70693 |
| *Adgrl4* | Adhesion G protein-coupled receptor L4 | 170757 |
| *Celsr1* | Cadherin, EGF LAG seven-pass G-type receptor 1 | 12614 |
| *Ephb2* | Eph receptor B2 | 13844 |
| *Efna2* | Ephrin A2 | 13637 |
| *Adgra2* | Adhesion G protein-coupled receptor A2 | 78560 |
| *Ephb3* | Eph receptor B3 | 1384 |

**Supplementary Table 5. Reagents, kits and antibodies.**

| **Reagents and kits** | **Company** | **Catalog Number** |
| --- | --- | --- |
| Formamide | EUROBIO | GHYFOR01.01 |
| Acetic anhydride | SIGMA | A6404 |
| Triethanolamine | SIGMA | T58300 |
| DMEM | Corning | 50-003-PCR |
| GlutaMAX^TM^ | Gibco | 35050061 |
| Penicillin-streptomycin | In Vitro | A-02 |
| Polyethylenimine | Polysciences | 23966-1 |
| Poli-L-lysine | Sigma-Aldrich | P2636-25MG |
| Paraformaldehyde | Electron Microscopy Sciences | 15714 |
| DAPI | Sigma-Aldrich | 10236276001 |
| TMB | Thermo Fisher Scientific | 00-202-3 |
| PBS | Corning | 55-031-PCR |
| Flag affinity resin | Thermo Fisher Scientific | 649493 |
| Coelenterazine 400a | GoldBIO.com | C-320 |
| BLOCK-iT Pol II miR RNAi Expression Vector Kits | Invitrogen | K4935-00 |
| HiFi assembly | New England Biolabs | E2621 |
| **Antibodies** | **Company** | **Catalog Number** |
| anti-DIG-POD | Roche | 11207733910 |
| anti-Fluorescein-POD | Roche | 11426346910 |
| anti-HA | BioLegend | 901514 |
| anti-Flag | Sigma-Aldrich | F7425-.2MG |
| anti-GFP | NovusBio | NB600-308 |
| anti-α tubulin | DSHB | 12G10 |
| anti-mouse IRDye800CW | LI-COR | C50924-02 |
| anti-rabbit IRDye680RD | LI-COR | C51104-08 |
| anti-mouse-HRP | MP Biomedicals | 0855550 |
| anti-rabbit-HRP | MP Biomedicals | 08670391 |
| anti-mouse-Alexa568 | Abcam | ab175472 |
| anti-mouse-Alexa488 | Abcam | ab150105 |

**SUPPLEMENTARY FIGURES**

**SUPPLEMENTARY FIGURE 1. *Adgrl1*, *Adgrl2* and *Adgrl3* mRNA expression in cortical and subcortical regions.  a-c.** Detection of *Adgrl1*, *Adgrl2* and *Adgrl3* (latrophilins 1,2 and 3) mRNA in cortical and subcortical areas using fluorescence in situ hybridization. Cx: cortex, Hipp: hippocampus, PO: posterior complex of the thalamus, LHA: lateral hypothalamic area. **d-e.** Functional validation of miRNAs against ADGRL1. Empty arrowhead: CTF with unknown post-translational modification. **f.** Genomic organization of *Adgrl1* alternatively-spliced exons and targeting of common exons by miRNAs used in this study.

**SUPPLEMENTARY FIGURE 2. Recombinant proteins schematics.** Schematic representation of the recombinant proteins used in this study. FS-like: follistatin-like, LDL-TM: transmembranal region of LDL receptor, EGF: epidermal growth factor, LNS: laminin-neurexin-sex hormone binding globulin, Glyc: glycosylation, PDZbm: PDZ binding motif, LEC: lectin domain, HBD: hormone binding domain, GAIN: GPCR Autoproteolytic Inducing domain, GPS: GPCR proteolysis site.

**SUPPLEMENTARY FIGURE 3. Hevin retention to NRX1α-expressing cells is reduced with calcium chelation.** DOLR assays in the presence of EGTA for NRX1α expressing cells. Fluorescence immunoblotting values for hevin are expressed as percentage of the condition without EGTA. Data represents mean values from technical replicates of a single experiment.

**SUPPLEMENTARY FIGURE 4. Hevin binds at the cell surface of ADGRL1 and NRX1α-expressing cells. a.** Confocal images from cell-surface labeling assays, showing hevin’s binding to NRX1α expressing cells. White arrowheads: spots for red and green channels strong overlapping. Yellow arrowheads: cells without detectable ADGRL1 and NRX1α expression. Scale bar 10 µm. **b-c.** Epifluorescence microscopy for the quantification of hevin total pixels in NRX1α expressing cells. Scale bar 100 µm. Data are represented as mean ± S.E.M of at least three independent experiments. Student's t-test, **p*<0.05.

**SUPPLEMENTARY FIGURE 5. Hevin impairs ADGRL1/NRX1β-mediated cell-cell adhesion. a.** Representative images from cell aggregation assays after 90 min of incubation. **b.** Quantification of the aggregation index for assays shown in a. **c.** Frequency of size distribution for aggregates shown in a. Data are represented as mean ± S.E.M of at least three independent experiments. Student's t-test, **p*<0.05.

**SUPPLEMENTARY FIGURE 6. Hevin modifies the relative steady-state presence of ADGRL1 fragments without affecting autoproteolysis. a-b.** Relative expression of NTF and CTF bands over GPS-intact receptor as shown in Figure 3b. **c-d.** Normalized abundance of NTF and CTF bands as shown in Figure 3b. Homekeeping protein α-tubulin was used for normalization. Data are represented as mean ± S.E.M of at least three independent experiments. Student's t-test, ****p*<0.001, *****p*<0.0001.

**SUPPLEMENTARY FIGURE 7.  Hevin’s detection increases in whole-cell extracts but decreases in extracellular media when co-transfected with ADGRL1.  a.** Immunoblot from whole-cell lysate for cells co-expressing hevin and ADGRL1. **b.** Quantification of fluorescence immunoblotting values for bands corresponding to hevin in a. **c.** Immunoblot from conditioned media for cells co-expressing hevin and ADGRL1. **d.** Quantification of fluorescence immunoblotting values for bands corresponding to hevin after 48 h of co-transfection with EV or ADGRL1 plasmids. Data represents mean values from technical replicates of a single experiment.

**SUPPLEMENTARY FIGURE 8.  ADGRL1 constitutively activates Gq and G11 biosensors.** ADGRL1 coupling to Gq and G11 was detected with the TRUPATH BRET-based biosensors. Normalized BRET^2^ ratio data are represented as mean ± S.E.M of at least three independent experiments. Two-way ANOVA, **p*<0.05, ***p*<0.01 ****p*<0.001, *****p*<0.0001. Student's t-test.

**SUPPLEMENTARY FIGURE 9. Hevin doesn’t affect TRUPATH biosensors in the absence of ADGRL1.** **a-c.** Normalized BRET^2^ ratio values for HEK293 cells (without ADGRL1 expression) after incubation with hevin. Data are represented as mean ± S.E.M of at least three independent experiments. Two-way ANOVA.

**SUPPLEMENTARY FIGURE 10. BRET-based G proteins biosensors unaffected by hevin exposure of ADGRL1-expressing cells. a-c.** Normalized BRET^2^ ratio values from G protein biosensors in ADGRL1-expressing cells which showed no changes after incubation with hevin. Data are represented as mean ± S.E.M of at least three independent experiments. Two-way ANOVA.
