## Supplementary figures and images for "Genetic liability underlying reward-related comorbidity in psychiatric disorders involves the coincident functions of autism-linked ADGRL1 and hevin"

### Supplementary Figure 1

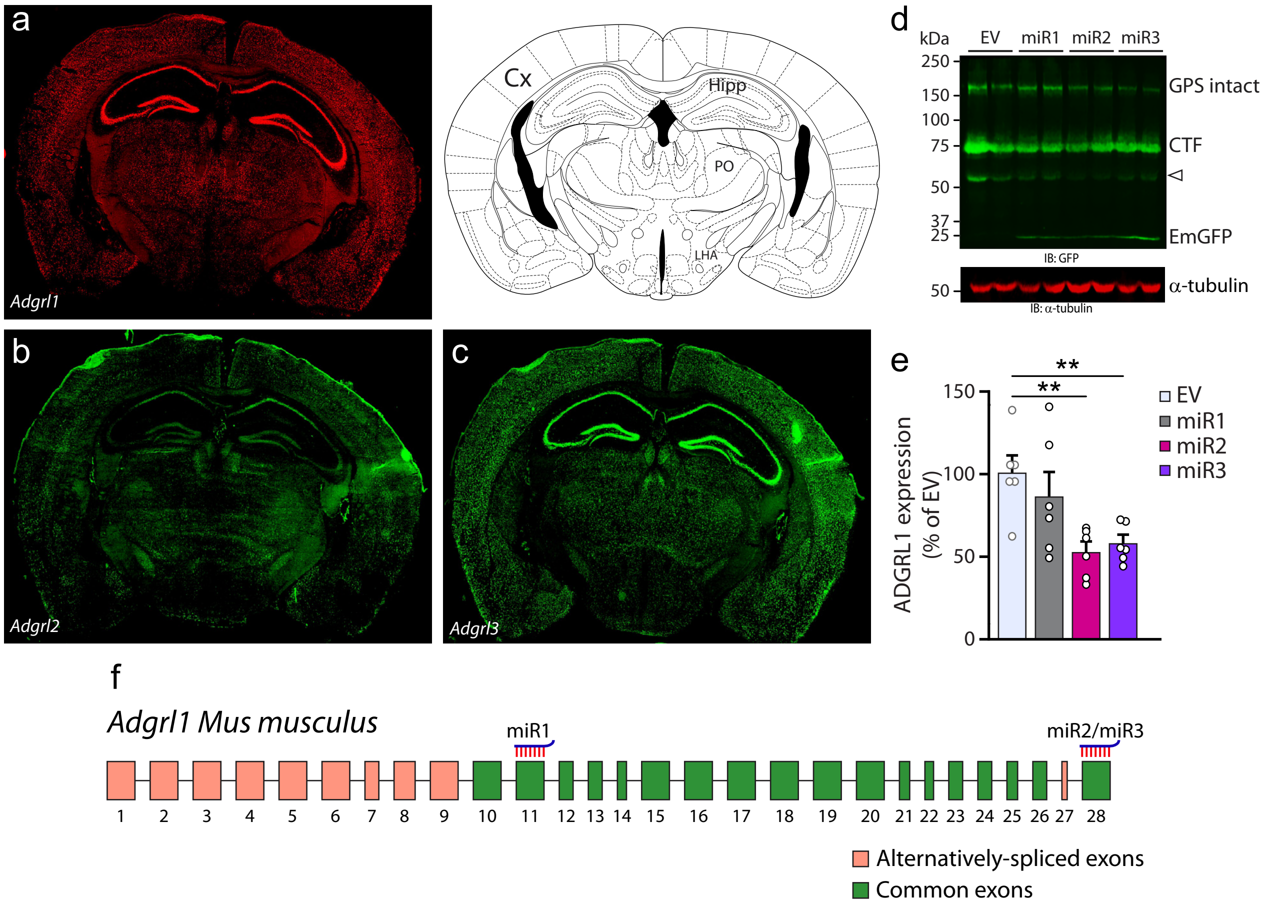

### Supplementary Figure 2

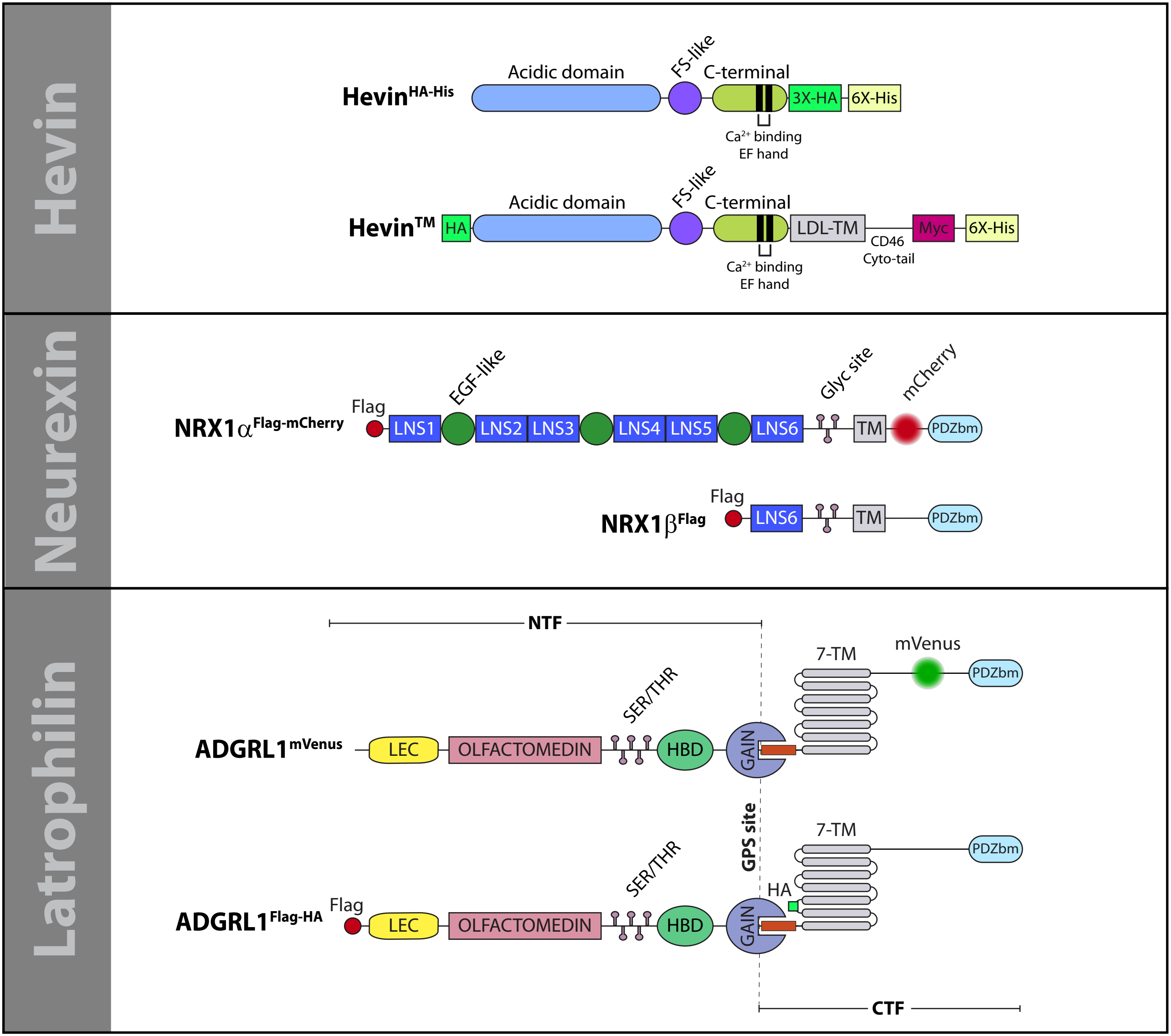

### Supplementary Figure 3

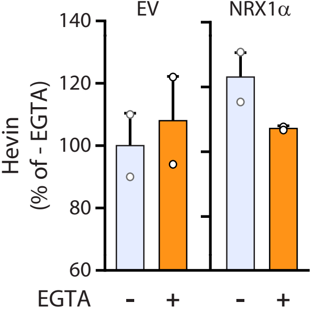


Suppl Fig 3

### Supplementary Figure 4

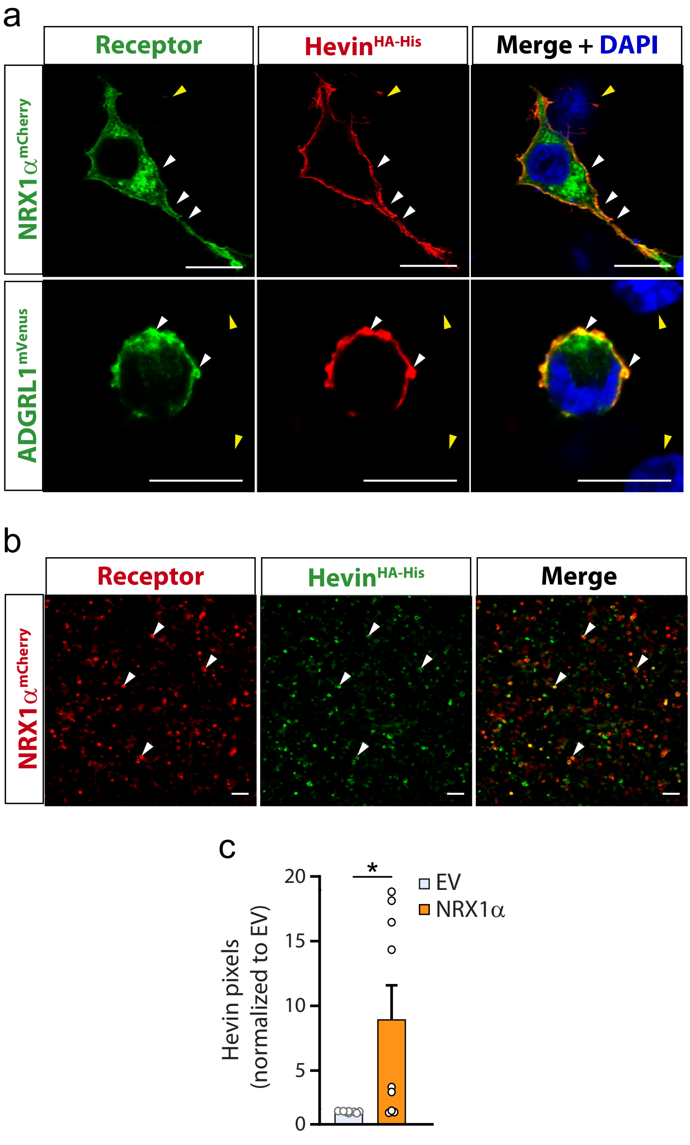

### Supplementary Figure 5

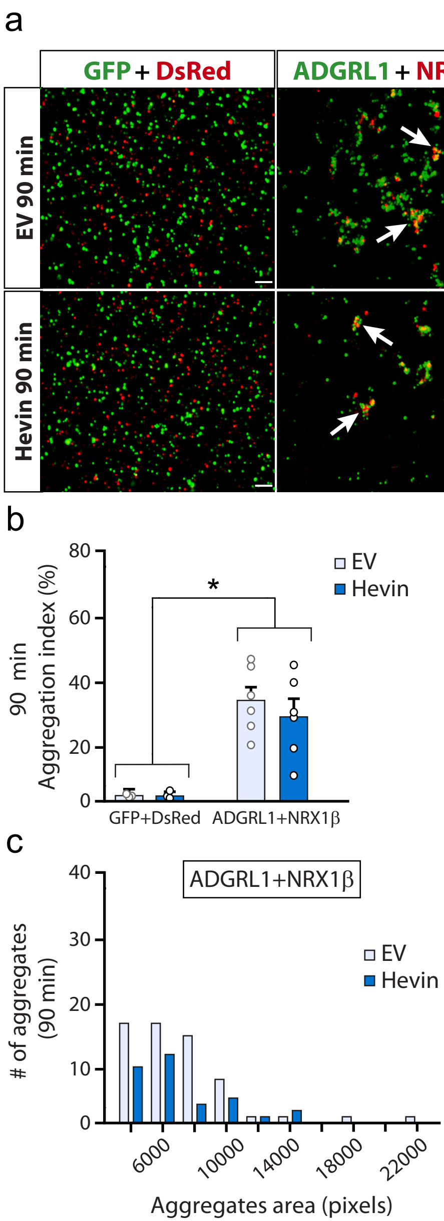

### Supplementary Figure 6

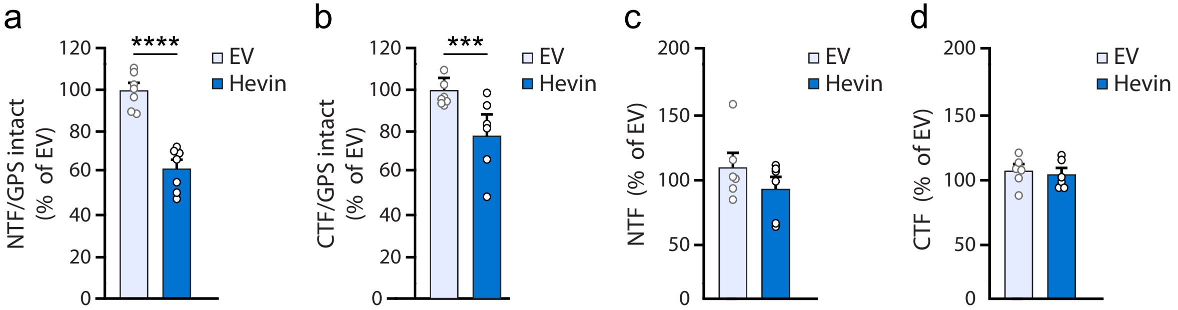

### Supplementary Figure 7

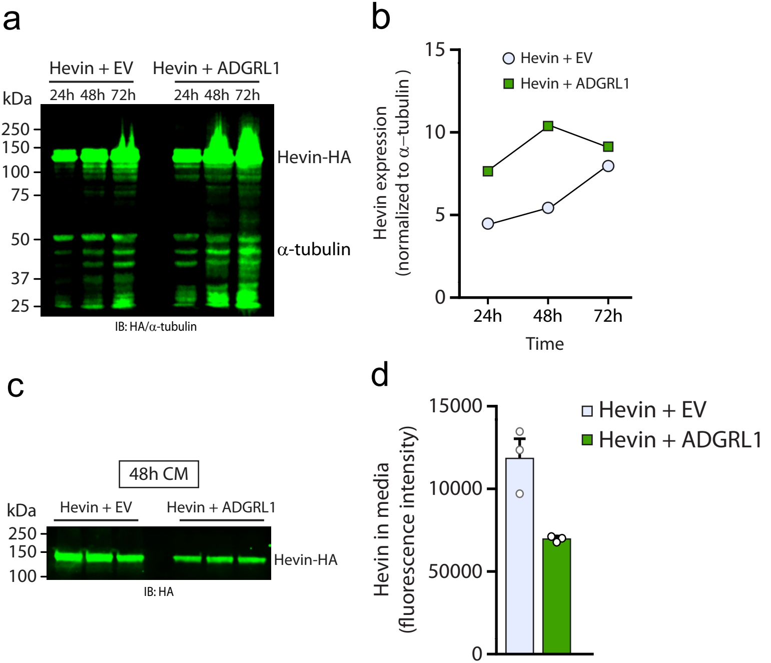

### Supplementary Figure 8

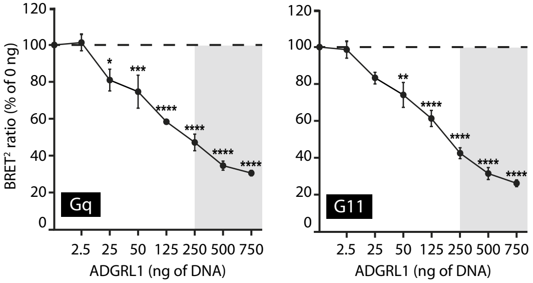

### Supplementary Figure 9

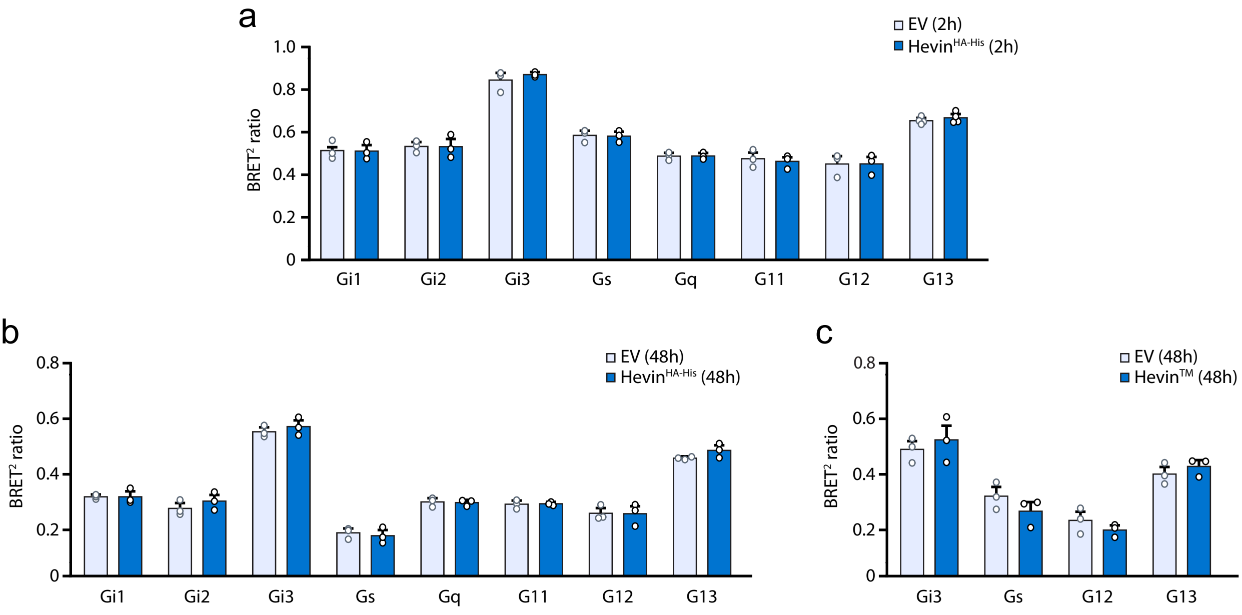

### Supplementary Figure 10

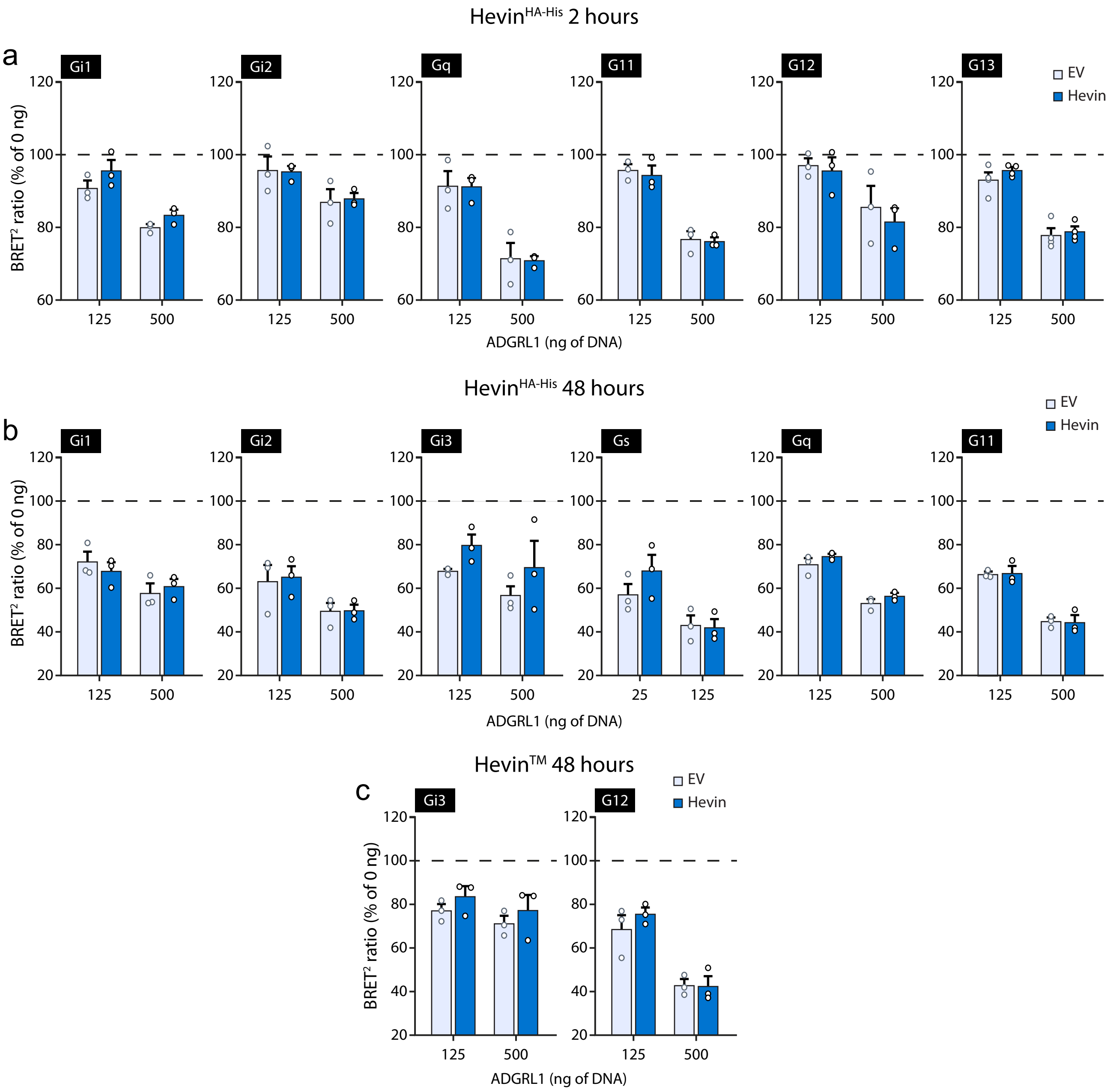
